# Multimodal on-axis platform for all-optical electrophysiology with near-infrared probes in human stem-cell-derived cardiomyocytes

**DOI:** 10.1101/269258

**Authors:** Aleksandra Klimas, Gloria Ortiz, Steven Boggess, Evan W. Miller, Emilia Entcheva

## Abstract

Combined optogenetic stimulation and optical imaging permits scalable, high-throughput probing of cellular electrophysiology and optimization of stem-cell derived excitable cells, such as neurons and muscle cells. We report a new “on-axis” configuration of OptoDyCE, our all-optical platform for studying human induced pluripotent stem-cell-derived cardiomyocytes (hiPSC-CMs) and other cell types, optically driven by Channelrhodopsin2 (ChR2). This solid-state system integrates optogenetic stimulation with temporally-multiplexed simultaneous recording of membrane voltage (V_m_) and intracellular calcium ([Ca^2+^]_i_) dynamics using a single photodetector. We demonstrate the capacity for combining multiple spectrally-compatible actuators and sensors, including newer high-performance near-infrared (NIR) voltage probes BeRST1 and Di-4-ANBDQBS, to record complex spatiotemporal responses of hiPSC-CMs to drugs in a high-throughput manner.

## 1. Introduction

Heart tissue is inherently dynamic, where a framework for both signal quantification and active interrogation is necessary to dissect the complex spatiotemporal phenomena underlying cardiac function [1]. Contraction of the heart is driven by the propagation of electrical waves. At the cellular level, the triggering event, action potential (AP), is determined by the balance of inward (depolarizing) and outward (repolarizing) ionic currents; the AP is closely followed by the increase in intracellular calcium concentration in the form of calcium transients (CTs), and ultimately calcium mediates mechanical contraction. To better understand electrical dysfunction, which can lead to complex electrical disturbances known as arrhythmias, optical mapping provides a means of contactless, high spatiotemporal recording of activity in cardiac tissue by employing fluorescent reporters. AP signals are obtained using fluorescent membrane voltage (V_m_) probes, such as the styryl dyes RH-237, Di-4-ANEPPS, and Di-8-ANEPPS, and intracellular calcium changes are captured by [Ca^2+^]_i-_sensitive probes, such as Fura-2, Fluo-4 and Rhod-4, which typically provide stronger signals compared to the commonly used styryl-based V_m_ probes [2–4]. When combined with spectrally-compatible optogenetic actuators, such as the genetically encoded depolarizing ion channel Channelrhodopsin2 (ChR2) [5] (**Fig. 1a,b**), dynamic events in cardiac electrophysiology can be interrogated in a fully contactless, high-throughput dynamic manner [6]. Dynamic space-time patterns of light can be imposed for control of excitation waves in cardiac tissue [7]. These optogenetic actuators can also be combined with genetically-encoded sensors, such as the genetically-encoded calcium indicators (GECI) R-GECO [8] (**Fig. 1b**) and GCaMP6f [9] (when a spectrally-compatible version of ChR2 is used), as well as voltage-sensitive fluorescent proteins (VSFP) such as VSFP3 [10] and QuasAr2 [11]. Newer red-shifted near-infrared (NIR) voltage-sensitive dyes, producing superior optical signals, such as Di-4-ANBDQBS [12] and BeRST1 [13], not only expand the palette of available sensors combinable with optogenetic actuators, but also enable simultaneous recording of V_m_ and [Ca^2+^]_i_ of optically interrogated samples when using green-excited [Ca^2+^]_i_ probes.

The traditional tools for probing cardiac electrophysiology, such as the planar patch clamp systems [14] and microelectrode arrays (MEAs) [15], rely on contact-based interrogation and often are limited to isolated/single cells and/or (non-cardiac) cell lines. In contrast, the contactless nature of all-optical platforms, makes them suitable for probing of electrical activity in two-and three-dimensional constructs of human induced-pluripotent stem cells (hiPSC-CMs), i.e. allows them to investigate electrical activity within the tissue context. Our previously described system OptoDyCE [6] exemplifies how optical mapping combined with optogenetics could be used to construct a low-cost, all-optical system for high-throughput cardiac electrophysiology. Here, we show the next generation of the OptoDyCE platform (**Fig. 1c**), where “on axis” design is used for optical stimulation and simultaneous V_m_ and [Ca^2+^]_i_ imaging in hiPSC-CMs onto a single detector with “strobed” LED illumination, gated to specific camera frames (**Fig. 1d**). The “on axis” configuration refers to the combined single optical path leading up to the sample for stimulation and for the multiparameter imaging. The resultant system is compact and fully compatible with standard multi-well plates (**Fig. 1e**).

In order to characterize and demonstrate the flexibility of the system, we compare multiple V_m_ and [Ca^2+^]_i_ probes in hiPSC-CMs. In addition to the more commonly used NIR V_m_ sensor Di-4-ANBDQBS, we show the first full characterization of a new NIR probe, BeRST1, in optogenetically paced hiPSC-CMs and demonstrate the compatibility of both sensors with ChR2 actuation as well as the [Ca^2+^]_i_ probes Rhod-4 and the genetically-encoded R-GECO. We assess the performance of the OptoDyCE platform and the utility of these probes for all-optical cardiac electrophysiology by quantifying spectral cross-talk due to the simultaneous use of multiple fluorescent probes and the stability of these probes under long-term, strobed illumination. To demonstrate the value of simultaneous dual imaging with high spatiotemporal resolution in the context of cardiotoxicity testing, we explore cell-level uncoupling between V_m_ and [Ca^2^+]i in a known proarrhythmic compound, azimilide, in both spontaneous and optically-paced conditions. By fully characterizing the presented all-optical platform, we show that when combined with hiPSC-CMs or other scalable cell types, OptoDyCE can provide a high-throughput means of standardization of protocols for electrophysiology testing across multiple sites [16–18], cell types [19], as well as multiple testing platforms [19–21] with a high number of replicates making high-throughput drug discovery, disease modeling, and personalized medicine realizable.

## 2. Materials and Methods

### 2.1 Human iPS-cardiomyocyte culture and gene delivery

Culture of hiPSC-CMs and adenoviral delivery of ChR2(H134R) was performed as described previously [6,22]. Briefly, frozen human iPSC-derived cardiomyocytes (iCell Cardiomyocytes^2^ ™, Cellular Dynamics International (CDI), Madison, WI) were thawed per the manufacturer’s instructions and plated on fibronectin coated wells in 384-well glass-bottom plates (P384-1.5H-N, Cellvis, Mountain View, CA; **Fig. 1e**) at the recommended plating density of 156,000 cells/cm^2^ (17,000 cells/well for a 384 well plate). After 5 days, adenoviral delivery of ChR2(H134R)-eYFP to the iPSC-CMs was performed in-dish at a viral dose of MOI 350. Transfection of R-GECO (Addgene #45494 (CMV-R-GECO1.2) developed by Robert Campbell) was performed in-dish after delivery of ChR2. Briefly, Lipofectamine 3000 (ThermoFisher, Waltham, MA), P3000 (ThermoFisher), CDI iCell Plating Medium, and the R-GECO plasmid were combined per manufacturer’s instructions and plasmid-containing solution remained in each well for at least 48 hours. For all samples, functional testing was performed 2 days after transfection and/or infection.

### 2.2 All-optical recording

Membrane voltage (V_m_) and intracellular calcium ([Ca^2+^]_i_) were administered as described previously [6]. All experiments were performed at room temperature in Tyrode’s solution containing the following (in mM): NaCl, 135; MgCl2, 1; KCl, 5.4; CaCl2, 1.33; NaH2PO4, 0.33; glucose, 5; and HEPES, 5 at pH 7.4. Optical recording of V_m_ was performed using the synthetic dyes BeRST1 [13] and Di-4-ANBDQBS (L. M. Loew), while [Ca^2+^]_i_ was reported optically with Rhod-4 or R-GECO. All probes were spectrally-compatible with ChR2. BeRST1, Di-4-ANBDQBS, and Rhod-4 were diluted in Tyrode’s solution to 1 μM, 35 μM, and 10 μM, respectively. Optical imaging was performed at >200 frames per second (fps) with 4×4 binning with a field of view of ~0.4 mm^2^ using NIS-Elements AR (Nikon Instruments; Melville, NY) on the OptoDyCE platform [6]. Recordings of spontaneous and paced activity were obtained, where 5 ms, 0.5Hz optical stimulation (470 nm) was provided using supra-threshold irradiances, as needed (in all cases < 1mW/mm^2^).

### 2.3 Dual imaging of voltage and calcium

A schematic of the protocol along with the light path of the optical system can be seen in **Fig.1**, similar to that described previously [6]. The optical system (**Fig. 1c**) was built around an inverted microscope (Nikon Eclipse TE-2000-U) fitted with a programmable x-y stage (OptiScan ES107; Prior Scientific; Rockland, MA) and automated z-focus (PS3H122 Motorized Focus; Prior Scientific), with illumination for actuation and sensing provided by a custom-built adaptor. Illumination for sensing was provided by red (sLED1) and green (sLED2) for voltage and calcium, respectively (Vm: 640 mW LED at 660nm; or [Ca2+]i: 350 mW LED at 530nm, both from Thorlabs); both LEDs were fitted with a bandpass filter (F_ex1_ (Vm): 655/40 nm; F_ex2_ ([Ca2+]i): 535/50 nm).The actuation LED (aLED) for ChR2 (650 mW at 470 nm, Thorlabs) was fitted with a 470/28 nm bandpass filter, F_actu_. The light paths for optical sensing and actuation were combined by a dichroic mirror (495 nm long-pass) and directed to the sample. Irradiances between 0.3 – 6 mW/mm^2^ for V_m_ and 0.05 – 1 mW/mm^2^ for [Ca^2+^]_i_ were selected based on maximum signal strength while minimizing oversaturation of pixels and minimizing effects on ChR2 activation. Irradiances were chosen based on cell plating density and filter choice, but were kept constant over the course of a single experiment. In order to achieve simultaneous imaging, temporal multiplexing was used, where the LEDs for exciting the V_m_ and [Ca^2+^]_i_ probes were gated to each camera frame. Collimation optics comprised of several lenses (L), and an objective lens (in this case 20x Nikon CFI Super Plan Fluor) was used to direct light to the sample. Emitted fluorescence was collected by a photodetector (iXon Ultra 897 EMCCD; Andor Technology Ltd., Belfast, UK). Illumination control was completely automated via software and TTL pulses (**Fig. 1d**).

### 2.4 Drug preparation

Azimilide was diluted in DMSO (Sigma Aldrich) to 1000x concentrations to achieve concentrations of 0.01 – 10 μM when diluted in Tyrode’s solution. After the sample plates were stained, the Tyrode’s wash solution was removed and replaced with Tyrode’s solution containing the compound. The plates were then returned to the incubator (37°C, 5% CO_2_) for 30 minutes prior to experiments to allow for to equilibration.

### 2.5 Data processing and analysis

Data was analyzed using software custom developed in MATLAB, [6] and the following endpoints were extracted: peak, percent change in fluorescence (ΔF/F%), and mean and standard deviation of baseline of V_m_ and [Ca^2+^]_i_. Signal-to-noise ratio (SNR) was calculated as ratio of peak signal to the standard deviation of the signal baseline. Pre-processing included baseline correction, removal of artifacts, temporal filtering using a Savitzky-Golay polynomial filter (second order, 3 frame window) and normalization.

## 3. Results

### 3.1 On-axis dual imaging system

The OptoDyCE system has been adapted to perform, for the first time, simultaneous dual imaging of voltage and calcium along with optogenetic stimulation using a single-detector, on-axis system (**Fig. 1c**). The choice of sensors and actuators is primarily limited by the spectral properties of the relevant chromophores (**Fig. 1a**). Currently, dye choice is dictated by the most popular blue-excited actuator ChR2. Many relevan-sensors capable of excitation by LEDs are excited at wavelengths longer than ChR2, including the green-excited [Ca^2+^]_i_ sensors Rhod-4 and R-GECO [8] and red-excited V_m_ sensors, Di-4-ANBDQBS [12] and BeRST1 [13]. Choice of these probes then determined the configuration of the optical system. The LEDs for exciting the sensors (sLED1, sLED2) and the actuator (aLED) along with bandpass filters used to narrow their spectra to prevent overlap (F_ex1_, F_ex2_, F_actu_) were selected based on the absorption spectra of chromophores. Likewise, this determined the selection of the dichroic mirrors (DM1, DM2, DM3) used to combine excitation sources and separate emitted photons before passing through the emission filters in front of the detector. Given the relatively low light levels required and the large selection of economically-priced light sources, a palette of probes can be used depending on spectral compatibility.

**Fig. 1.**
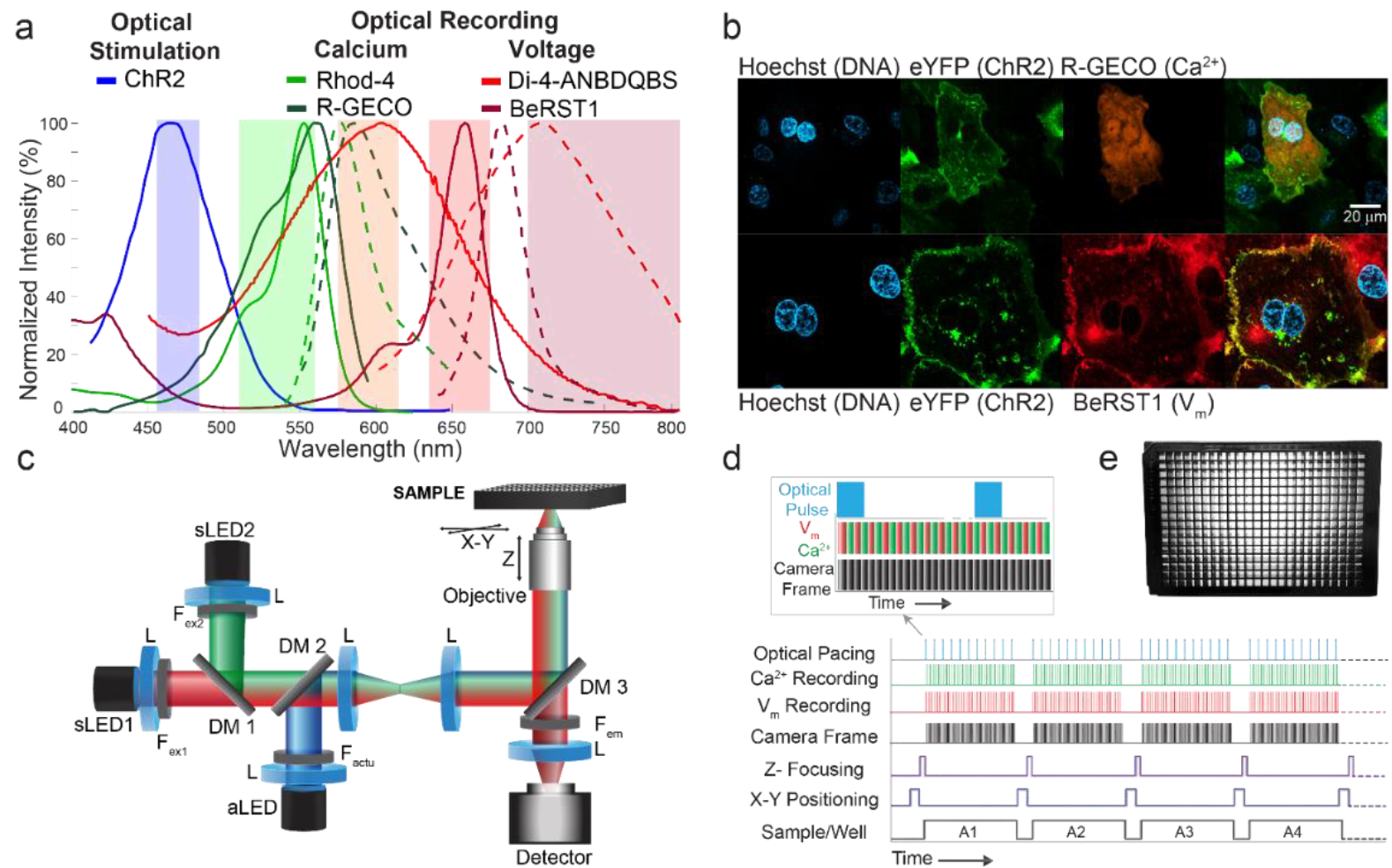
The OptoDyCE platform for “on axis” multimodal all-optical cardiac electrophysiology. (a) Spectrally-compatible synthetic and genetically-encoded fluorescent reporters of V_m_ and [Ca^2+^]_i_ can be combined with optogenetic actuators to perform all-optical interrogation of hiPSC-CMs. Fluorescent images (b) of hiPSC-CMs treated with the nuclear stain Hoechst (blue) show successful expression of both eYFP-tagged ChR2 (green) with R-GECO (orange), a genetically encoded [Ca^2+^]_i_ sensor, while the V_m_ dye BeRST1 (red) shows strong membrane localization. Scale bar is 20 μm. OptoDyCE (c) can readily perform simultaneous imaging of V_m_ and [Ca^2+^]_i_ using a single detector through ‘strobing’ the sensing LEDs (sLED1 and 2), which are temporally multiplexed or gated by the camera (d). Due to the contactless nature of all-optical interrogation, this approach allows for automated recording of standard multi-well plates (e).

### 3.2 Compatibility of V_m_ and [Ca^2+^]_i_ probes for dual imaging: cross-talk quantification

Simultaneous imaging using V_m_ and [Ca^2+^]_i_ probes, including the genetically encoded [Ca^2+^]_i_ sensor R-GECO, reliably produces signals (**Fig. 2a,c**) with minimal crosstalk (**Fig. 2 b,d**), demonstrating the applicability of the platform for multiparameter assessment of function. Crosstalk of Di-4-ANBDQBS into the [Ca^2+^]_i_ records is present at this scale in the form of minor motion artifacts (**Fig. 2a,c**), however the SNR and ΔF/F(%) are small compared to that of Rhod-4. Crosstalk of the remaining probes is negligible, with detector counts close to that of the dark counts of the EMCCD camera. Dual-strobing of dual-stained samples produced signals comparable in morphology to that of single-probe stained samples, both in terms of detector counts and ΔF/F(%). Dual-strobing versus single strobing does produce some crosstalk, where the strong Rhod-4 is seen to bleed into on the V_m_ channel (no green illumination) in single-stained samples (**Fig. 2c,d**), most likely the result of the non-zero decay time of the green LED, which can be resolved by reducing the exposure time of the illumination LED. An interesting observation of increase in ΔF/F(%) and SNR for combined BeRST1 and Rhod-4 imaging (**Fig. 2c,d**) does not appear to be due to LED bleed-through, as the signal increase is observed under both single and dual strobing conditions and is specific to BeRST1 (not present for Di-4-ANBDQBS), and warrants further investigation. We argue that SNR provides a more informative metric for the quality of the optical signals compared to ΔF/F(%), as seen when comparing V_m_ and [Ca^2+^]_i_ probes, where the voltage probes may produce lower ΔF/F(%) but the obtained signals have very high SNRs and require minimal post-processing.

**Fig. 2.**
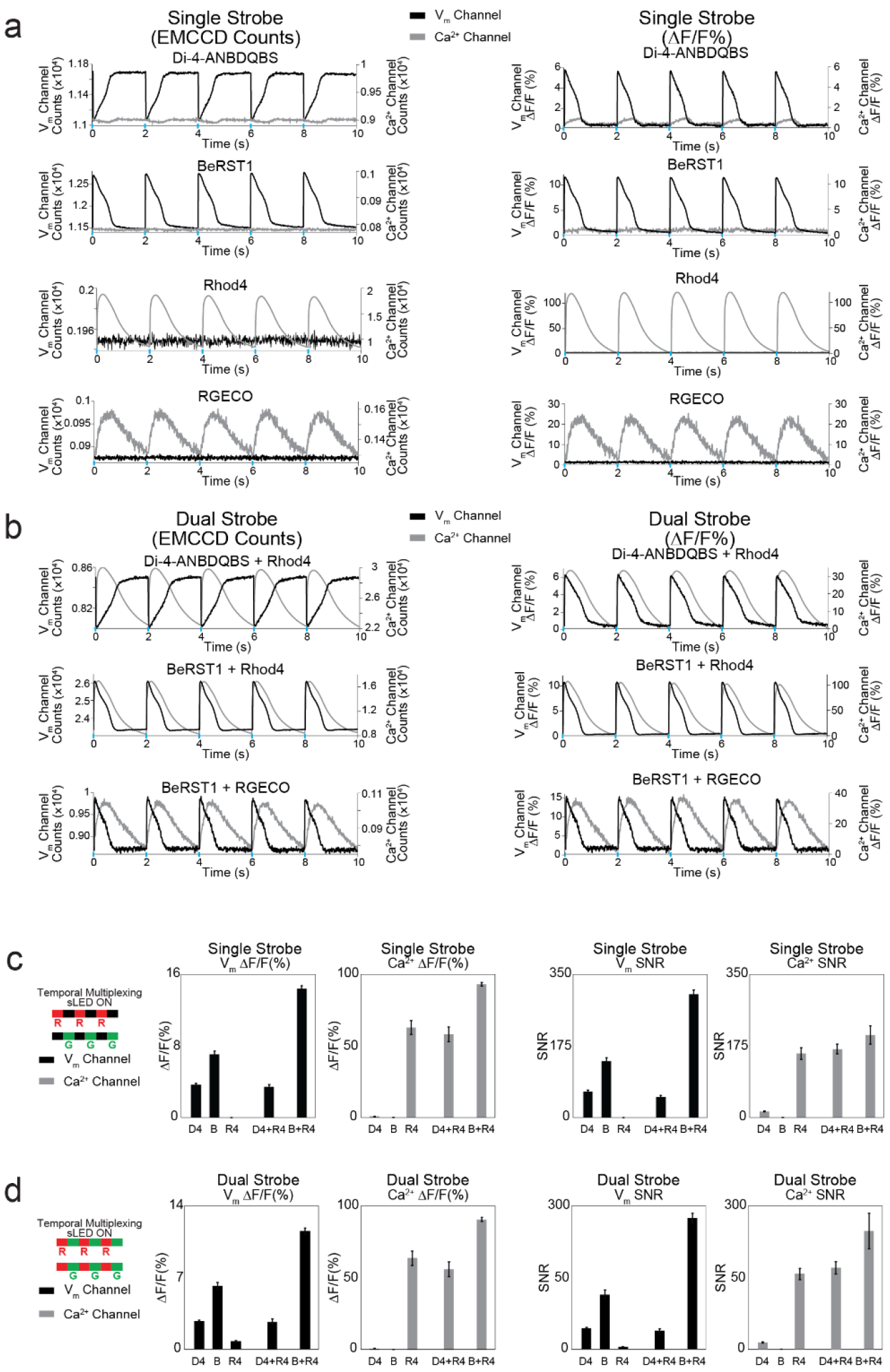
Investigation of compatibility of V_m_ and [Ca^2+^]_i_ probes and cross-talk. OptoDyCE can simultaneously measure V_m_ and [Ca^2+^]_i_ using Di-4-ANBDQBS (D4) or BeRST1 (B) combined with Rhod-4 (R4) or RGECO (RG) under strobed illumination of hiPSC-CMs, optically paced at 0.5Hz. V_m_ and [Ca^2+^]_i_ channel denote illumination under the excitation source (sLED) of the respective probes (red and green). Signals are given in terms of EMCCD detector counts (left) and ΔF/F(%) (right) for both single (a) and dual (b) strobe conditions. Single strobe illumination of single-probe samples (a, left) show counts close to the detector dark counts for B, R4, and RG while red-excited D4 is seen to produce a motion-artifact signal on the [Ca^2+^]_i_ channel (green illumination). Comparison of ΔF/F(%) to SNR for both single-strobe and dual-strobe illumination (N = 5,6) (c,d) show that even if present, the SNR of the minor-crosstalk is much smaller than that of the desired signal for the respective channel. Irradiances were kept the same for a particular wavelength (dye type) within a plate, but were adjusted between plates to account for differences in cell plating and dye loading and to optimize SNR; red LED: 1.4 to 2.5 mW/mm^2^, green LED 0.46 to 0.53 mW/mm^2^.

### 3.3 Quantification of photobleaching and suitability of V_m_ and [Ca^2+^]_i_, probes for longterm imaging

To determine the suitability of the synthetic V_m_ and [Ca^2+^]_i_ probes for long term measurements, 5 minute continuous recordings under strobed illumination were obtained in hiPSC-CM samples, optically paced at 0.5Hz (**Fig. 3**). Bleaching occurs more strongly with Rhod-4 compared to the voltage probes (**Fig. 3a,b**), with a greater drop in ΔF/F(%) and SNR observed with Rhod-4 when comparing the first 20 seconds and last 20 seconds of the recording (**Fig. 3b**). BeRST1 did not exhibit any photobleaching within this 5-minute record, and was generally stable for hours of experimentation. Di-4-ANBDQBS had more baseline drift, most likely due to debris entering and exiting the field of view at this microscale; this was also present to a lesser extent in BeRST1 samples which appeared to be less taken up by cellular debris. Even in the presence of bleaching, no major changes in signal morphology were observed with any of the three probes, and in all three cases SNR remained of high quality (**Fig. 3c**). None of the probes used in this study was found to result in any phototoxicity-induced changes in AP/CT morphology within the typical 1-3 hour recording window.

**Fig. 3.**
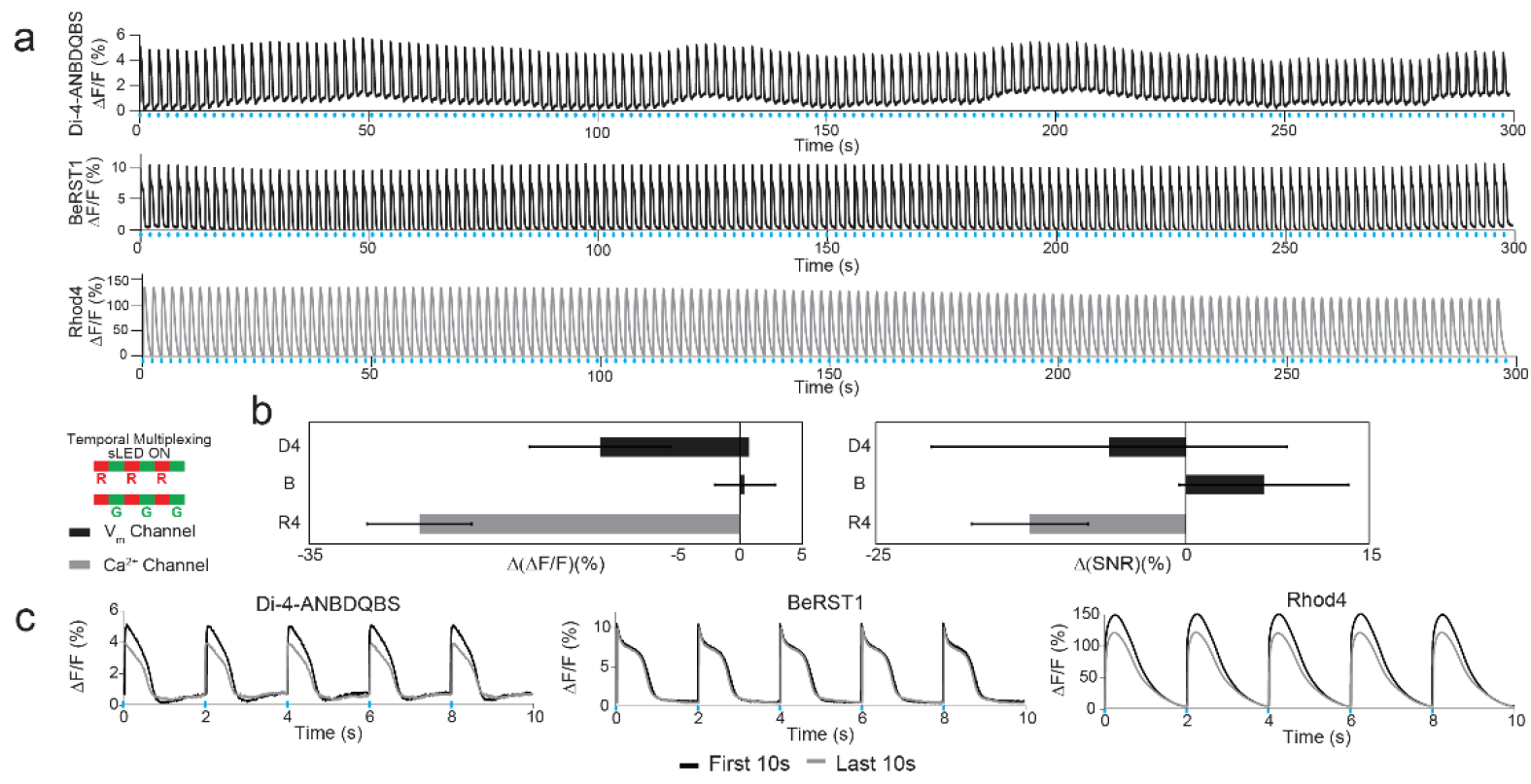
Stability and photobleaching of V_m_ and [Ca^2+^]_i_ probes. Strobed illumination for 5 minutes combined with optical pacing exhibits minor photobleaching for di-4-ANBDQBS (D4) and Rhod4 (R4), but not BeRST (B) (a) with stronger bleaching observed in Rhod4 (R4) when comparing the percent change between the first and last 20 seconds of the recording for both ΔF/F(%) (given as Δ(ΔF/F)) and SNR (given as Δ(SNR)) (N=4) (b). Additionally, no major changes in AP/CT morphology are observed after 5 minutes of recording (c).

### 3.4 Spatiotemporal imaging

To demonstrate the utility of simultaneous dual imaging using OptoDyCE, hiPSC-CMs were treated with azimilide, a known pro-arrhythmic compound. Spontaneous and 0.5Hz optical pacing recordings of hiPSC-CMs stained with BeRST1 and Rhod-4 were obtained (**Fig. 4**). Although the azimilide-treated samples (**Error! Reference source not found.b**) showed AP and CT prolongation (known pro-arrhythmic markers) compared to DMSO control (**Error! Reference source not found.a**), no decoupling of voltage and calcium was observed in the global signal, averaged over the whole field of view. However, signals from sufficiently small ROIs at the single-cell scale show such decoupling of V_m_ and [Ca^2+^]_i_ (another pro-arrhythmic feature) in azimilide-treated samples, where abnormal activity in the spontaneous [Ca^2+^]_i_ signal is not reflected in the V_m_ signal. This highlights the need for simultaneous multiparameter measurements and for spatially-resolved imaging. Pacing can be effective in reducing this decoupling and in suppressing abnormal calcium release events, yet the drug-induced prolongation limits the rates at which the system remains stable and responsive.

**Fig. 4.**
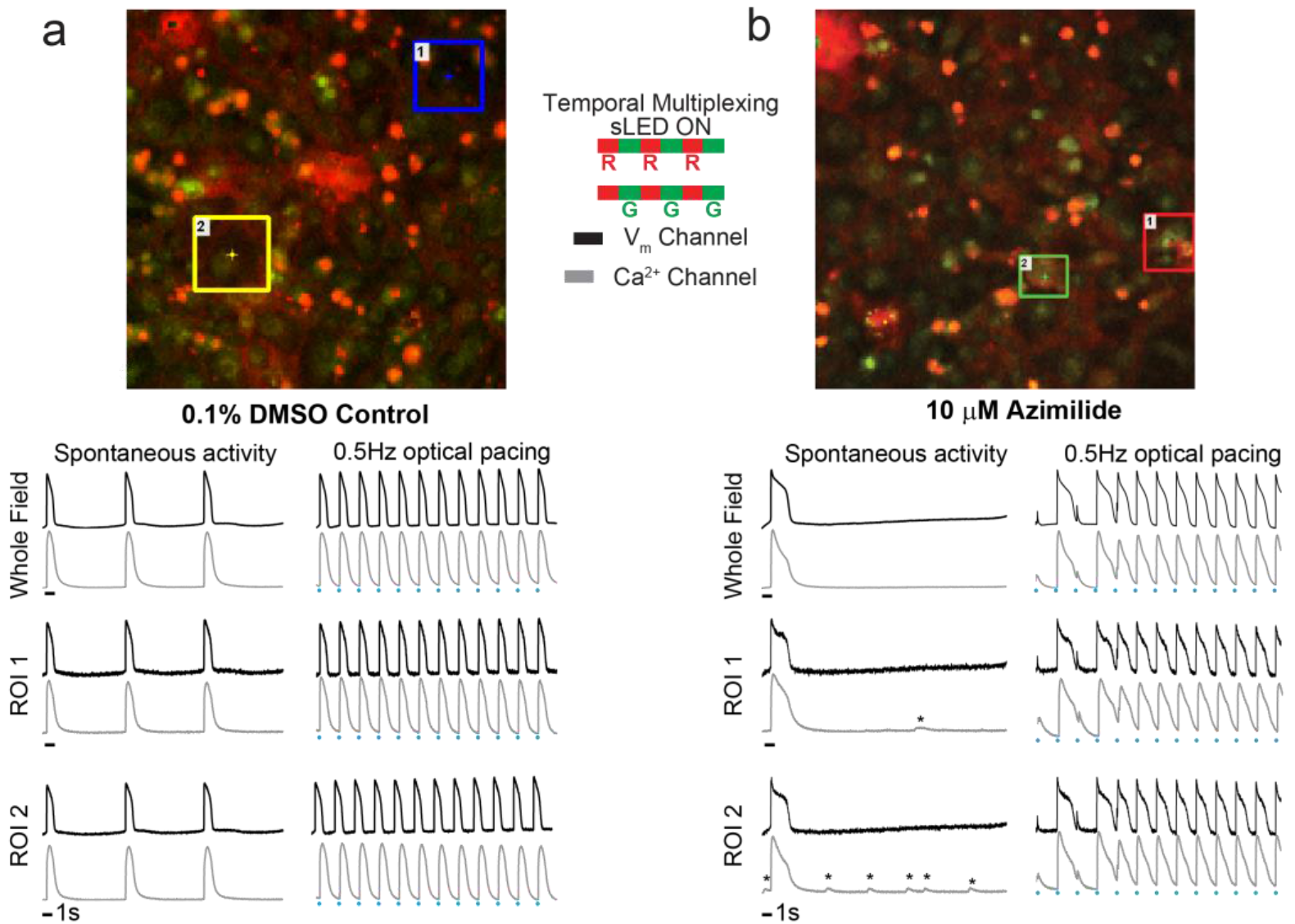
Dual imaging of V_m_ and [Ca^2+^]_i_ probes to capture sub-cellular events for pro-arrhythmia prediction. hiPSC-CM samples stained with BeRST1 and Rhod-4 were treated with 0.1% DMSO (a) or 10 μM azimilide (b), a known proarrhythmic compound. Averaging over the whole field, azimilide is seen to prolong both the spontaneous and paced action potential and calcium transient, a known marker of pro-arrhythmic risk. Signals from single-cell ROIs show decoupling of V_m_ and [Ca^2+^]_i_ in azimilide treated samples (^*^asterisks), another indicator of pro-arrhythmic risk, not detectable in the global signal.

### 3.5 Effects of excitation irradiance on SNR

Although ΔF/F% is the most commonly reported quantification of signal quality in optical mapping, SNR plays a larger role in signal processing and feature detection. In this study with the presented platform, all probes showed excellent SNR. Even for very low light levels and low concentrations of staining, the optical signals were usable without filtering. The three probes characterized here were capable of SNR of 100 or more. Varying incident irradiance of the green LED for Rhod-4 excitation (**Fig. 5a**) shows that SNR only scales directly with irradiance until 0.3 mW/mm^2^, where oversaturation of pixels results in decreasing SNR. It should be noted that the LED was set to only 30% maximum power (0.53 mW/mm^2^) before the whole detector area was oversaturated for these samples. Although ChR2 kinetics are affected by green (520 nm) wavelengths [22,23], peak SNR of Rhod-4 appears to occur at relatively low irradiances and thus excitation of the probe should not engage ChR2. Both Di-4-ANBDQBS and BeRST1 show similar profiles for irradiance (**Fig. 5b,c**). SNR for BeRST1 plateaus around 2.8 mW/mm^2^, while Di-4-ANBDQBS decreases in SNR at high LED irradiances > 5.45 mW/mm^2^ due to pixel oversaturation in areas with dye-containing cellular debris. For both V_m_ probes, oversaturation of the whole detector area did not occur before the LED power was at maximum (6.6 mW/mm^2^). For most recordings, excitation irradiances ranged from 0.46 – 0.53 mW/mm^2^ for Rhod-4 and 1.4 – 2.5 mW/mm^2^ for Di-4-ANBDQBS and BeRST1, as seen in (**Fig. 5**) and (**Fig. 2c**).

**Fig. 5.**
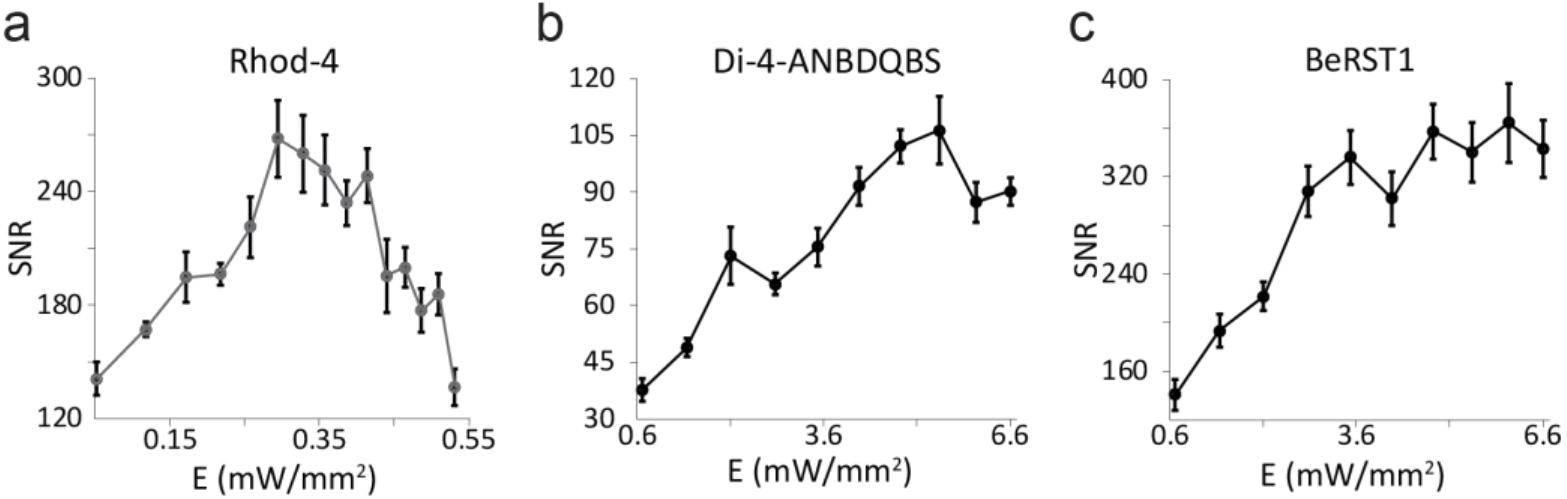
Effects of excitation irradiance on SNR of V_m_ and [Ca^2+^]_i_ probes. SNR for Rhod-4 (a), Di-4-ANBDQBS (b), and BeRST1 (c) were calculated for a single hiPSC-CM sample optically paced at 0.5Hz.

## 4. Discussion

Here we demonstrate that OptoDyCE can be adapted to perform simultaneous dual imaging of V_m_ and [Ca^2+^]_i_ combined with optogenetic actuation using ChR2 to interrogate complex spatiotemporal phenomena in hiPSC-CMs. Although dual imaging approaches using a single detector and two LED sources have been employed previously, these systems relied on an “off-axis” configuration, where the multiple illumination and detection paths do not overlap [24–26]. While dual “on-axis” approaches have been performed, they typically rely on two-detector configurations [27–32]. Considering that the photodetector (camera) is the most expensive component in these all-optical systems, the “on axis” temporal multiplexing approach presented here offers a particularly attractive solution. Additionally, none of the previously published techniques have demonstrated combined simultaneous measurements of V_m_ and [Ca^2+^]_i_ with optogenetic stimulation. Dual imaging using combined genetically-encoded sensors and actuators has been used to perform drug screening in hiPSC-CMs [33], however it required two cameras and two light sources for readout, and the actuators and sensors were not co-expressed in the same cells. Approaches using genetically-encoded probes and actuators are of interest in long-term monitoring of chronic drug treatment over hours and days, such as kinase-effecting drugs, where synthetic probes may not be ideal. OptoDyCE employs simple off-the-shelf optical components and uses temporal multiplexing of optical sensing by high-speed gating of fluorescent illumination not possible with traditional sources [34]. Imaging of complex phenomena can be performed using a single detector thus achieving a cost-effective yet powerful configuration capable of acquiring high-content information. The “on-axis” aspect of the system makes it amenable to miniaturization and integration with micro-endoscopic systems for all-optical *in vivo* applications [35].

As novel imaging technologies continue to be developed for use in cardiac electrophysiology, quality metrics for both the imaging system and the optical reporters must be established. Although optical mapping commonly uses ΔF/F% to determine the quality of a fluorescent reporter, we suggest that SNR provides a better metric of signal detectability and signal quality, which is more commonly used in the context of high-speed imaging. Given the need for recording speeds >200 fps (and thus short exposure times), signals with small (<10%) dynamic changes can be more susceptible to noise (e.g. detector and photon noise), requiring temporal and spatial filtering, which further degrades the measured signals [36]. When variability in the signal baseline is not taken into account, as with ΔF/F%, signal quality can be incorrectly represented, as seen when comparing V_m_ and [Ca^2+^]_i_ probes (**Fig. 2c,d**). Additionally, we observe that ΔF/F% varies more than SNR across samples, most likely due to variability in staining and illumination intensity. Probes producing higher SNRs lessen the need for filtering signals, which both maintains signal fidelity and reduces computational requirements. It also reduces the strict dependence on expensive, high-sensitivity cameras and large numerical aperture (NA) optics. Regardless, both NIR probes used in this study (Di-4-ANBDQBS and BeRST1) show high quality by both metrics, and can yield impressive SNRs, comparable to those for traditional [Ca^2+^]_i_ probes. It is important to note that in their current formulations, BeRST1 required two orders of magnitude lower concentration compared to Di-4-ANBDQBS (1μM compared to 35μM) to produce the signals presented here. However, further improvement of the SNR for Di-4-ANBDQBS and removal of the baseline drift may be possible by leveraging its capability as an excitation-ratiometric probe [12]. When coupled with suitable photodetectors, sensitive in the NIR, these newer voltage sensors represent an important contribution to optical electrophysiology.

The OptoDyCE platform is compatible with both synthetic and genetically-encoded sensors and allows for short-term and long-term all-optical control of samples with optogenetic actuators. In particular, the contactless nature makes the platform suitable for high-throughput measurements of hiPSC-CMs, allowing dissection of the V_m_ - [Ca^2+^]_i_ dynamics. Although in this study we have only demonstrated OptoDyCE as a tool for measuring optically reported V_m_ and [Ca^2+^]_i_ in hiPSC-CMs, the platform is suitable for recording any optically-measurable parameter (e.g. organelle-based ion concentrations, pH, contraction, etc.) and is impartial to the studied cell-type, allowing for the interrogation of compatible samples of different geometries, including 2D monolayers and 3D tissue constructs [6]. As new actuators and sensors (both synthetic and genetically encoded) become available, they can be easily screened with and then integrated into the platform. The ability to provide dynamic pacing and high-throughput recording on an all-optical “on-axis” platform, easily adaptable to changing technologies, provides an economical tool for better understanding of cardiac electrophysiology *in vitro* and *in vivo.* In the current *in vitro* context, it not only provides a means to improve drug discovery and cardiotoxicity testing, but also offers a cost-effective means of phenotypic testing of hiPSC-CMs in order to improve iPSC technologies for use in drug discovery and development, disease modeling, and personalized medicine.

